# Searching for sympatric speciation in the genomic era

**DOI:** 10.1101/367623

**Authors:** Emilie J Richards, Maria R. Servedio, Christopher H Martin

**Affiliations:** Department of Biology, University of North Carolina at Chapel Hill, Chapel Hill NC

**Keywords:** sympatric speciation, genomics, gene flow, introgression, selection, cichlid

## Abstract

Sympatric speciation illustrates how natural and sexual selection may create new species in isolation without geographic barriers. However, recent genomic reanalyses of classic examples of sympatric speciation have revealed complex histories of secondary gene flow. Thus, there is a need to revisit how to connect the diverse theoretical models of sympatric speciation and their predictions to empirical case studies in the face of widespread gene flow. We summarize theoretical differences between sympatric speciation and speciation-with-gene-flow models and propose genomic analyses for distinguishing which models apply to case studies based on the timing and function of adaptive introgression. Investigating whether secondary gene flow contributed to reproductive isolation is necessary to test whether predictions of theory are ultimately borne out in nature.

## What is sympatric speciation?

As Mayr famously quipped, sympatric speciation is like the Lernean Hydra: “which grew two new heads whenever one of its heads was cut off” (p. 451; [1]). The latest incarnation of this phenomenon has occurred over the past decade: sympatric speciation now means two different things to different research groups.

In the classic definition, sympatric speciation represents the most extreme endpoint on the divergence with gene flow continuum: panmictic gene flow and no initial divergence at the start of speciation [2–4]. In the context of theoretical speciation models, this classic process of sympatric speciation is the most difficult because the starting conditions involve no pre-existing divergence, potentially tied with physical linkage, among loci involved in reproductive isolation. Instead, **linkage disequilibrium** (see Glossary) must build up through time within a population through the action of disruptive natural selection and strong assortative mating by ecotype, despite the countervailing eroding force of recombination [5–8].

Recently, the definition of sympatric speciation has been expanded to focus more on biogeographical context [9], in which the speciation process is defined as sympatric a) as long as diverging populations are within ‘cruising range’ of each other and b) regardless of whether secondary gene flow provided alleles contributing to reproductive isolating barriers in sympatry. Cruising range provides a practical empirical definition of gene flow between diverging sympatric populations, allowing for some geographic or microallopatric population structure (e.g. [10,11]). However, allowing for secondary gene flow of alleles contributing some or all of the reproductive isolation between sympatric populations expands the definition of sympatric speciation to include both hard and easy processes under one umbrella. Theoretical models show that speciation is much easier from starting conditions that involve some level of initial divergence and/or restricted gene flow, for example, if the alleles necessary for reproductive isolation first become physically linked in allopatry [12].

These two definitions, the classic theoretical population genetic and the recent biogeographic, reflect different perspectives on the value of studying sympatric speciation. The biogeographic definition, with its broader range of starting conditions that are easier to verify in nature, increases, perhaps vastly, the number of empirical speciation events that could be categorized as examples of sympatric speciation. This definition values the frequency of sympatric diverging populations in nature compared to allopatric speciation, as an estimate of the overall importance of “sympatry” in contributing to biodiversity. The classic definition, with its narrow set of starting conditions that remain challenging to verify in nature, finds value in studying the difficult process of sympatric speciation itself. Namely it values the theoretical possibility of creating new species solely through the power of divergent selection alone, regardless of whether this process is common in nature. This opinion piece focuses on the types of questions that genomic data now allow us to ask to improve the search for examples of the latter process and understand the range of speciation mechanisms found in nature from among those shown to be plausible in theory.

Under the biogeographical definition of sympatric speciation, there is little difference in terms of the speciation mechanisms involved in scenarios that start with initial panmixia (e.g. classic sympatric speciation) versus those that start with some geographic or microallopatric population structure (e.g. [10,11]). This contrasts sharply with the rich theoretical literature differentiating models of sympatric speciation from other models of speciation with gene flow. Indeed, theory teaches us that the classic process of sympatric speciation (without the aid of secondary gene flow contributing to reproductive isolation) is uniquely and notoriously difficult [13], in part because quite specific conditions of resource availability (e.g., [6,14]), mating traits and preferences (e.g., [15,16]), and search costs (e.g., [17]) must be met for it to occur. Some claim that the effort to discern the exact geographic scenario and initial conditions of speciation would be better spent on finding loci involved in reproductive isolation (i.e. ‘barrier loci’ [18,19]). This is an important first step and we can glean something about the process of speciation from gene annotations of barrier loci and linkage architecture. However, understanding whether any one particular locus or potential mechanism was necessary for speciation often requires placing genomic discoveries in the context of speciation models that compare the importance of such processes, models whose outcomes are highly dependent on the initial conditions before sympatric divergence.

## Different mechanistic processes underlying divergence in sympatry

Regardless of definition, it is necessary to distinguish among different sympatric divergence processes to understand which classes of speciation models and predictions apply to specific case studies. We here distinguish different scenarios (Fig. 1) that will result in two sister species in sympatry based on whether **secondary gene flow** aided in population divergence: 1) classic sympatric speciation without gene flow; 2) sympatric speciation in the presence of a) neutral secondary gene flow or b) after differential sorting of an ancestral **hybrid swarm**. In the latter case, it is important to distinguish whether the ancestral hybrid swarm population achieved panmixia before later divergence (i.e. sympatric divergence); otherwise, differential sorting of haplotypes within the hybrid swarm is better described by secondary contact speciation with gene flow models rather than sympatric speciation models. 3) Speciation may also be aided by secondary gene flow that a) triggers initial sympatric divergence or b) increases divergence after initial divergence in sympatry becomes stalled, an outcome of many sympatric speciation models without sufficiently strong disruptive selection [20–23]. Finally, 4) secondary contact after a period of allopatry between two populations can result in coexistence or reinforcement, if there is not collapse into a single admixed population [12,24–26] or extinction of one or both populations. We consider scenarios 1 and 2a to be examples of classic sympatric speciation models, the ‘hard’ process, whereas scenarios 3 and 4 would be examples of speciation aided by secondary gene flow, a much easier process in theory. Interestingly, hybrid swarm scenarios (2b) exist in a gray area, since substantial initial gene flow from multiple sources may increase ecological or preference variation within a population that is sufficient to trigger later sympatric divergence, even without segregating inversions or genetic incompatibilities [27–29]. So far, we know of no examples of scenario 1 within any case study of sympatric sister species examined using genomic tools; even long diverged species show some evidence of **introgression** in their past (e.g. [30]). In contrast, sympatric speciation with neutral gene flow (Scenario 2a, and conditionally Scenario 2b) and speciation aided by gene flow (Scenarios 3 and 4) frequently appear to operate concurrently even within a single sympatric adaptive radiation (e.g. [31–33]).

**Figure 1:**
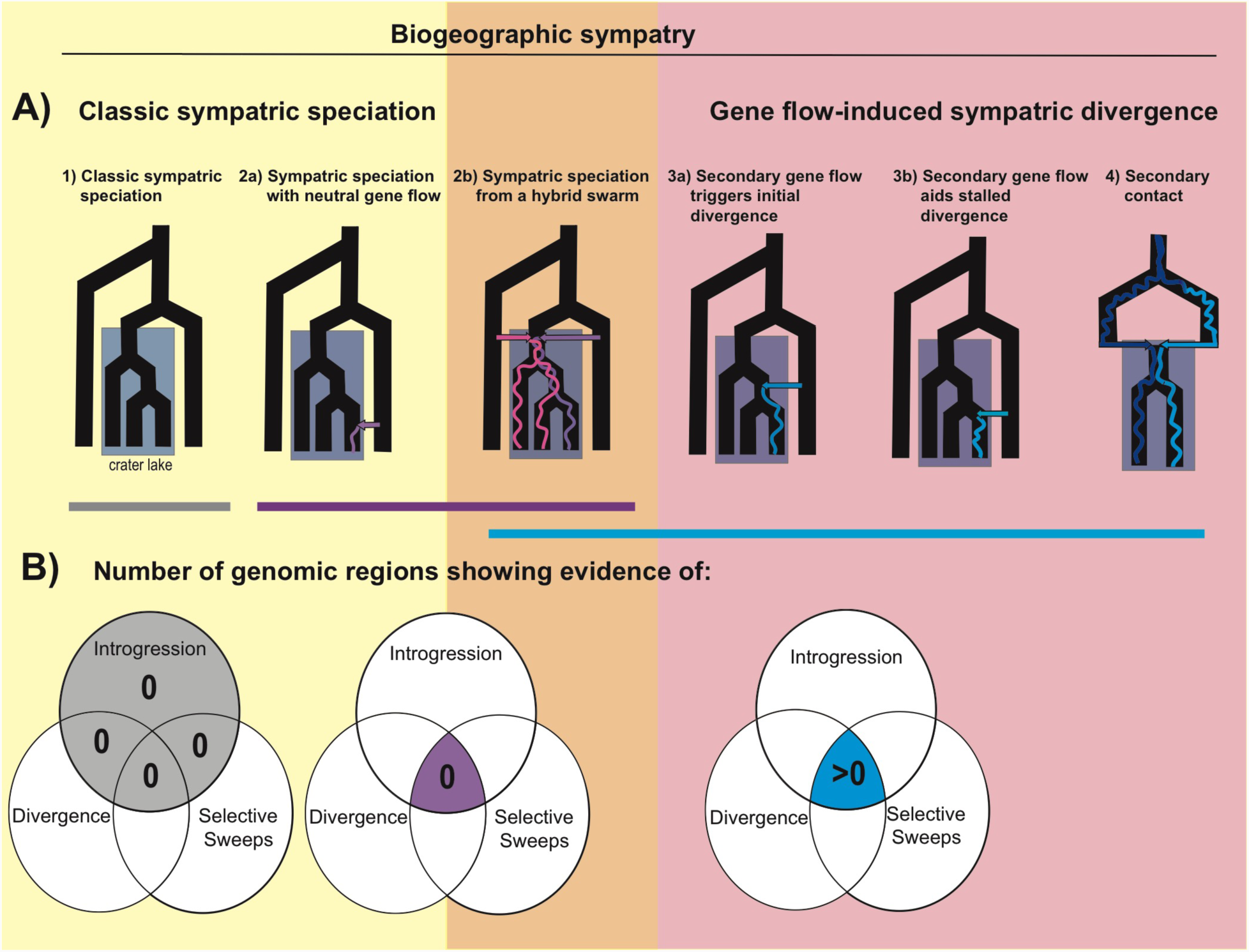
Genomic signatures of sympatric speciation and speciation with gene flow. Speciation scenarios are grouped into classic sympatric speciation scenarios (yellow box; harder process in theoretical models) and other divergence scenarios which can also occur under biogeographic sympatry (red box; easier processes, which we refer to broadly as sympatric divergence here). Speciation from a hybrid swarm (orange box) can fall under either class of scenarios and additional information is necessary to determine what category of speciation models best describe this process. **A)** The timing of gene flow relative to divergence can be used to distinguish between speciation scenarios. The colored arrows represent gene flow events and the colored lines within the tree are simplified representations of a signature of introgression from that gene flow event into the sympatric species. **B)** Venn diagrams illustrating the number of genomic windows across the entire genome expected to have overlapping signatures of introgression (e.g. *f*_*d*_ outliers), genetic divergence (e.g. *F*_*st*_ and *D*_*xy*_ outliers), and selective sweeps (e.g. SweeD) for each speciation scenario (e.g. see [115]). The highlighted sections of the Venn diagrams indicate the key signature that can be used to distinguish between the scenarios. The scenarios that are expected to leave very similar signatures of overlap are grouped by the bars colored with their respective Venn diagram.

It is important to distinguish these scenarios because theoretical models predict that sympatric divergence unaided by any form of secondary gene flow is substantially more difficult than other speciation with gene flow scenarios (Box 1). Gene flow throughout the speciation process allows recombination to break down linkage disequilibrium among alleles associated with ecological divergence and assortative mating. There are actually three different types of sympatric speciation models to consider: the most difficult process involves independently segregating loci for ecotype, female preferences, and male traits within the population, whereas sympatric divergence is much easier if any of these three types of traits are combined, such as assortative mating based on phenotype matching instead of separate loci for preference and traits [34,35] or “magic” traits (such as assortative mating based on microhabitat preference; [6,7,36]). Sympatric speciation by sexual selection alone is also theoretically possible (albeit considered highly unlikely) if there is substantial preference variation either initially within the population or through secondary gene flow [16,27].

Any form of linkage disequilibrium among ecological and mate choice loci formed in allopatry, whether due to physical linkage, selection, or drift, can tend to shift the initial starting conditions of panmixia in favor of sympatric divergence [24]. However, linkage disequilibrium without physical linkage subsides within a relatively small number of generations after secondary sympatry and thus may not allow sufficient time for the evolution of assortative mating within the population. In contrast, pre-existing physical linkage among ecological loci has been shown to increase the probability of divergence, especially when it captures already divergent alleles, as is more likely after a period of divergence in allopatry before secondary contact [37,38]. Similarly, physical linkage can cause preference and trait alleles to mimic phenotype matching, although even tight linkage can break down over long timescales (shown in a model with population structure: [39]). Segregating inversions in the ancestral population are now well-known empirical examples of physical linkage promoting divergence in sympatry [37,40]. Sympatric divergence is also limited by many other restrictive conditions regarding the costs of female choosiness and strengths of disruptive selection and assortative mating.

Despite extensive searches for examples of sympatric speciation in the wild, there are few convincing case studies due to the difficulty of ruling out historical allopatric scenarios (see below) and the new difficulty of ruling out a role of introgression in speciation. Furthermore, the role of magic traits or matching vs. preference/trait mechanisms is not fully understood in any existing case study. Thus, we still have very limited empirical tests of an extensive theoretical literature and diverse competing models of the notoriously difficult process of sympatric speciation [34,35,41,42].

## The classic problem of sympatric speciation

There are four traditional criteria for demonstrating classic sympatric speciation (Scenario 1): 1) sister species which are reproductively isolated, 2) form a **monophyletic** group, 3) largely overlap in ranges, and 4) have biogeographic and evolutionary histories that make periods of allopatric divergence highly unlikely [43]. Very few case studies have been able to meet these rigorous criteria despite intensive searches [4,43]. This has led to the prominent status of crater lake cichlid radiations as some of the best examples of sympatric speciation in the wild due to the uniform shape of isolated volcanic lakes which convincingly rule out phases of allopatry due to water level changes (Box 2; [44]).

The monophyly criterion assumes that monophyly arises only when a single ancestral population underlies the present-day daughter species. This is typically met by inferring a single phylogeny from one or more loci. This single point-estimate view of evolutionary history is problematic because it obscures the presence of non-bifurcating relationships among organisms (e.g. sister species which derived ancestry from multiple source populations due to extensive gene flow or hybrid speciation) and the real variation in evolutionary histories among genes across the genome itself (e.g. [45]). Few regions of the genome may initially contribute to reproductive isolation resulting in a heterogeneous genomic landscape of differentiation among incipient species [46], a pattern now extensively supported across case studies (e.g. [47–49]). Therefore, monophyletic relationships are consistent with, but not exclusive to a scenario of sympatric speciation. Examining heterogeneous evolutionary histories across regions relevant to speciation is thus crucial for understanding the processes and conditions under which sympatric divergence can occur.

## The ‘new’ problem of sympatric speciation

While genomics has increased our ability to resolve evolutionary relationships among organisms, it has also revealed more complex evolutionary histories of multiple colonizations and extensive secondary gene flow in nearly all examples of sympatric speciation that have been examined with genomic data so far ([50–59]; e.g. to our knowledge Lord Howe Island palms and indigobirds have not yet been directly examined for secondary gene flow with an outgroup). Indeed, only a handful of genes may directly contribute to the speciation process whereas the rest of the genome is porous to gene flow while reproductive isolation is incomplete [46,60]. Examples of sympatric speciation without secondary gene flow (Scenario 1) are now even rarer after applying modern genomic tools to search for introgression. Instead, it is still possible that sympatric speciation occurs in the face of secondary gene flow in nearly all these examples (Scenario 2; [56]). Importantly, most evidence of secondary gene flow impacting putative examples of sympatric speciation comes from genome-wide tests of introgression from outgroup lineages that do not look at how that secondary gene flow has impacted gene flow between diverging populations in sympatry (e.g. [53,54]). It is also possible in cases studies of sympatric speciation that involve radiations of species, that secondary gene flow impacted some, but not all of the divergence events such that some species within a radiation may better represent classic sympatric speciation scenarios than others. Therefore, introgression detected at the genome-wide level from lineages outside the speciation event tells us that secondary gene flow has occurred, but little about the divergence process among incipient sympatric species and how that gene flow shaped the process of speciation.

The challenge of understanding the hard process of classic sympatric speciation in the genomic era is establishing or rejecting a functional role for the ubiquitous secondary gene flow present during the speciation process, in effect ruling out scenarios 3 and 4 in favor of scenario 2 (Fig. 1). Even if signatures of secondary gene flow are detected, speciation could still have occurred solely via mechanisms of sympatric speciation, in the classical sense, if that secondary gene flow did not play a causal role in divergence (Scenario 2a, conditionally Scenario 2b). In contrast, secondary gene flow could play a causal role if it introduced novel genetic variation or physically linked alleles (e.g. a segregating inversion) that promoted speciation before the start of divergence, through mechanisms such as inflating variance through the creation of a hybrid swarm (Scenario 2b; ([28,29,32,61]), adaptive introgression (Scenario 3; [62–64]), **transgressive segregation** (Scenarios 2-3; [65,66]), or hybrid speciation ([67]). Here we propose and discuss genomic analyses that may help to establish or reject a functional role of secondary gene flow in the speciation process (Fig. 1). This is necessary to identify putative cases of the classic sympatric speciation process when gene flow appears to be nearly universal in the wild, particularly among sympatric diverging populations.

## Searching for sympatric speciation in the genomic era

Although genome-wide analyses of introgression provide a starting point, ultimately consideration of the time of arrival and functional role of each introgressed region within extant sympatric sister species pairs will be necessary to distinguish between the classic ‘hard’ process of sympatric speciation in which incidental gene flow does not contribute to reproductive isolating barriers (Scenario 2a) or speciation aided by secondary gene flow (Scenario 3; e.g. segregating inversions [68,69] or balancing selection on regions containing multiple barrier loci [70,71]). We suggest four major types of genomic analyses to address questions about the role of secondary gene flow and identify sympatric speciation with gene flow: analyses to1) estimate the timing of introgression into sympatric sister species relative to their divergence time, 2) infer the presence and timing of selective sweeps within sympatric sister species, 3) annotate candidate adaptive introgression regions for functional elements or trait associations that may be relevant to speciation, and 4) if closely related non-speciating outgroups are available, confirm the lack of selective sweeps of these regions in outgroups. These analyses are by no means trivial, but recently developed methods have made it possible to start addressing such challenging questions.

The geographic context and breeding locations of populations are also needed to sort classic sympatric divergence from parapatric or microallopatric divergence scenarios. In particular, sympatric taxa which segregate in different microhabitats, such as different depths or host types, exhibit a type of automatic magic trait (i.e. ‘mate where you eat’) and belong to the easiest class of sympatric speciation models if these microallopatric traits diverged in sympatry [68,72]. Combining geographic and natural history context with population genetic statistics will aid in distinguishing where case studies fall along the speciation with gene flow continuum and whether the starting conditions of panmixia in sympatric speciation models will apply (Fig. 1).

### 1) Is the observed secondary gene flow concurrent with divergence times?

Estimating the duration of gene flow and the timing of introgression into the sister species from an outgroup relative to the timing of divergence between sympatric sister species will help distinguish between scenarios of sympatric speciation, speciation with gene flow, and secondary contact. If populations diverged in sympatry independent of any concurrent secondary gene flow (Scenario 2), we might expect to see weak concordance of the timing of gene flow with divergence times among species, for example discrete gene flow events that date well before or after divergence times among species (Fig 1A). In the case of both discrete gene flow events surrounding divergence time estimates or continuous gene flow from the time of colonization to the present, more information about function and selection on regions introgressed near the time of speciation will be needed. Increasingly sophisticated approaches for detecting fine-scale patterns of introgression and inferring the timing and duration of gene flow from genomic data are becoming available (Box 3).

### 2) Are any of the introgressed regions under selective sweeps and does the timing of these sweeps align with species divergence time?

We can use information about selective sweeps of introgressed variation to further characterize the role of secondary gene flow in sympatric divergence. When an allele is selectively favored in a population, positive selection may cause it to increase in frequency and form a localized selective sweep of reduced genetic variation surrounding the adaptive variant [73]. Such regions of high differentiation in recently diverged species are often targeted as candidates for speciation genes, although other processes not directly associated with speciation can lead to similar patterns of high heterogeneity in differentiation across a genome (reviewed in [19,74,75]). If speciation was recent or ongoing, there may be strong signatures of a selective sweep for particular haplotypes in at least one of the sister species for regions involved in the divergence process (Fig. 1B). If secondary gene flow was neutral with respect to speciation, we may find no signatures of selective sweeps in those introgressed regions.

Importantly, a sweep of the same introgressed region in both sympatric sister species may be interpreted as adaptation to the same new environment, which may not contribute to reproductive isolation between the pair (dependent on their respective genetic backgrounds; e.g. [76,77]). However, this pattern is also consistent with the sweep of a region contributing to a ‘one-allele’ mechanism of mate choice [5,34,35], such as increased female choosiness in both sympatric sister species (e.g. [78]), which *would* contribute to reproductive isolation. Thus, selective sweeps of an introgressed region in both sympatric sister species do not rule out its role in aiding the speciation process.

Alternatively, if selective sweeps are detected, the timing of selective sweeps can give indirect evidence about their role in speciation. If the timing of introgression predates the timing of the selective sweep, it is challenging to infer the importance of an introgressed region for speciation because linkage disequilibrium among loci relevant to speciation may take time to build up. However, the absence of selective sweeps or introgression until long after species divergence would suggest that introgression was not relevant to speciation (Scenario 2a).

### 3) Is there support for a causal role of secondary gene flow based on functional genetic analyses of variants in the region?

Another potential source of evidence for the functional importance of gene flow can come from genome-wide association studies (GWAS) between variants in introgressed regions and traits involved in ecological or sexual isolation between sister species. The conservation of sequences within introgressed regions across taxa may also provide strong evidence of a functional role (e.g. PhastCons [79]). However, many complex traits are driven by a large number of variants of small effect and ruling out a functional role for gene flow from gene annotations is difficult (e.g. see the omnigenic model; [80]). Finally, and most powerfully, genome editing and gene expression reporter systems are increasingly tractable in non-model systems (e.g. [81,82]). This is ultimately an asymmetric problem: finding evidence that an introgressed region may have contributed to reproductive isolation is far easier than demonstrating that no introgressed regions contributed to reproductive isolation in any way [56]. Finding evidence for sympatric speciation in the wild is now the difficult problem of functional genetic analyses of introgressed regions.

### 4) Are there similar patterns of selection or divergence in the introgessed regions in closely related outgroup populations?

Thorough investigation of these same regions in outgroups to the sympatric species gives added power to distinguish whether secondary gene flow aided sympatric divergence. If non-diversifying, closely related species exist in similar environments and haven’t diversified in a similar manner but share signatures of selective sweeps in the same regions, then the observed introgression may have been neutral relative to speciation, e.g. due to adaptations to shared changes in climate or pathogens or shared regions of reduced recombination or increased background selection. Similarly, several studies comparing genomic landscapes of differentiation across closely related taxa have found that high differentiation observed in the same genomic regions across taxa reflects the action of linked selection across low-recombination regions rather than selection against gene flow at barrier loci [83–86].

## Concluding Remarks

Sympatric speciation remains among the most controversial evolutionary processes, beloved by theorists and long sought after by empiricists. While evidence of divergence under the biogeographic definition of sympatry is mounting using traditional genetic criteria of monophyly [4], genomic data has now revealed the ubiquity of secondary gene flow and introgression in these many of these examples. Future fine-scale investigations of introgression will likely continue to paint a complex picture of the role of secondary gene flow in speciation. Ruling out a role for secondary gene flow in speciation and discerning which putative cases studies meet the classic definition of sympatric speciation in the wild will be a formidable task, yet a worthwhile one in its revelation of the sheer power of divergent selection to create species in nature.

Nearly all existing case studies of sympatric speciation involve some form of automatic magic trait, such as assortative mating by habitat [68,87,88], along a depth gradient [52], or environment-induced phenology shifts [89]. We think an outstanding remaining question is whether the classic ‘hard’ process of sympatric speciation occurs in nature without the aid of some form of magic trait, as originally demonstrated to be possible in theory [6]. The highly polygenic and multi-dimensional nature of adaptation and mate choice suggests that an ‘all-of-the-above’ speciation scenario containing a mix of preference/trait, magic trait, and phenotype matching, each spread across a wide distribution of allelic effect sizes with varying times of arrival, will be the norm in nature. In contrast, although numerous and diverse, most speciation models continue to address these mechanisms in a piecemeal fashion with an assumption of large effect alleles. It remains unclear how different mechanisms, effect sizes, and times of arrival will interact and compete within a single model.

## *Box 1.* Why do we care whether speciation is sympatric?

Inferences from theoretical models predict that, under a scenario of speciation with gene flow (Scenario 3), introgression can make the process of speciation much easier in three ways. First, by introducing additional variation in ecological traits into the population, introgression could potentially facilitate a branching process due to competition for resources (although we are not aware of a model that assesses this precise situation, it can be inferred from the dynamics of [6]). Second, introgression of novel alleles for mating preferences may provide a boost in preference variation that could be an important trigger to aid the evolution of assortative mating under a preference/trait mechanism, which requires preference variation to be large ([16,27]). For example, we found evidence of secondary gene flow of olfactory alleles shortly before the rapid divergence of a Cameroon cichlid radiation in Lake Ejagham, which may have boosted preference variation [55]. Third, secondary sympatry may lead to increased linkage disequilibrium between assortative mating and ecological loci or among ecological loci. It seems logical that this might facilitate sympatric speciation as this metric is often described as progress along the speciation continuum. However, initial linkage disequilibrium has been shown not to matter much in at least some scenarios [5] because without physical linkage, linkage disequilibrium breaks down quickly. However, physical linkage may enable these alleles to remain in association for a sufficient time for assortative mating to evolve within the population (e.g., [39]). Initial linkage disequilibrium may also increase the probability of allelic capture by an inversion or for selection for new mutations within the inversion that may affect both ecology and assortment [37]. Finally, higher linkage disequilibrium among ecological loci may in some cases increase the probability of sympatric divergence, but this is in effect similar to varying effect sizes of alleles at ecological loci (e.g. many small effect alleles within a region resemble a large-effect locus [90]). These predictions could also apply to sympatric radiations. For example, some classic sympatric speciation models (e.g. [6]) can yield many more that just two species if left to run for more generations [14,22].

The fundamental difference between sympatric speciation and speciation with gene flow, including secondary contact scenarios, lies in the fact that very often multiple equilibrium states exist in speciation models, such that loss of divergence and maintenance of divergence in the presence of gene flow are both possible outcomes, depending on the starting conditions of a population (this is nicely illustrated for one measure of divergence by [24], Fig. I). In such cases, speciation is much more easily reached from starting conditions that match those of two populations that have diverged largely in allopatry due to the large amount of allelic variation or pre-existing phenotypic bimodality and assortative mating. Even for scenarios of speciation with gene flow that are much easier, such as geographic separation between two incipient species that are undergoing gene flow, differentiation is much more difficult to reach or maintain from an initially homogeneous population than from an initially differentiated one [91,92].

**Fig I.**
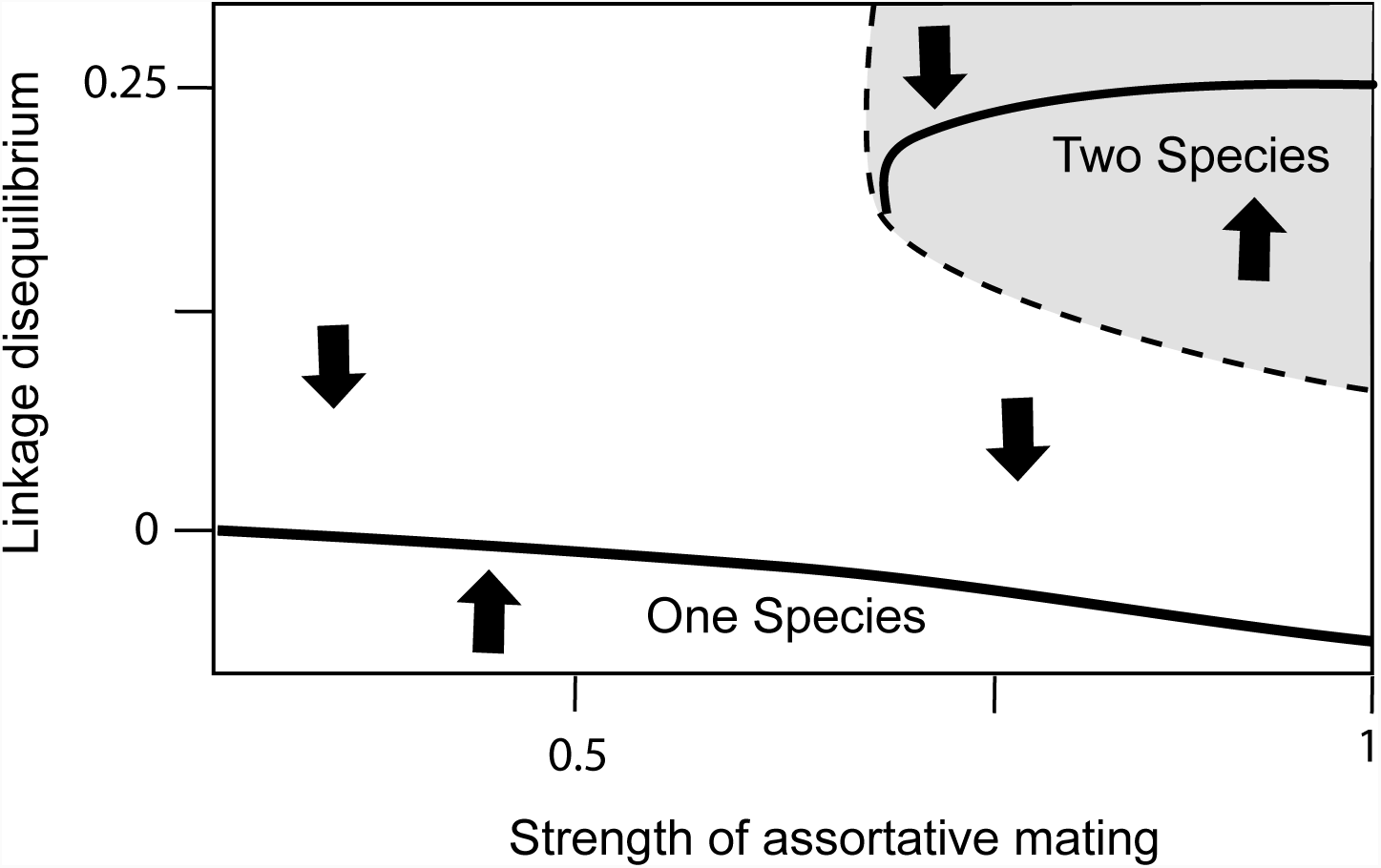
Two equilibrium cases exist for the linkage disequilibrium (LD), a proxy for differentiation into two distinct “species” in this proof-of-concept model, that can be maintained between two loci that are under disruptive selection and determine assortative mating. With little initial LD, the one-species equilibrium is likely to be reached even when the intensity of assortment is high. When LD in the traits is initially large, as can be the case if there is initially divergence in allopatry, the two-species equilibrium can be reached instead. Modified from [24].

## *Box 2.* Evidence for sympatric speciation from crater lake cichlid radiations

There are relatively few volcanic chains of crater lakes containing fishes in the tropics, notably found in Cameroon, Nicaragua, Tanzania, Uganda, Madagascar, and Papua New Guinea [52,93,94]. Although sympatric radiations of endemic fishes are known from other isolated saline, alkali, postglacial, and ancient lakes, only four lineages of cichlids have radiated in the world’s crater lakes (Fig. II). The most diverse radiation is Barombi Mbo, Cameroon with eleven endemic cichlid species, followed by Lake Bermin, Cameroon with nine [95]. Nicaraguan crater lakes reach up to five species [96], the East African craters never exceed two sympatric species [52,57], and Madagascar’s crater lakes contain a single endemic cichlid [94]. It remains unknown why regional and lineage diversity varies so greatly because there appears to be no relationship between the occurrence of endemic cichlid radiations and crater lake size or age (up to approximately 5 km diameter and 2 million years old) [97].

**Fig. II.**
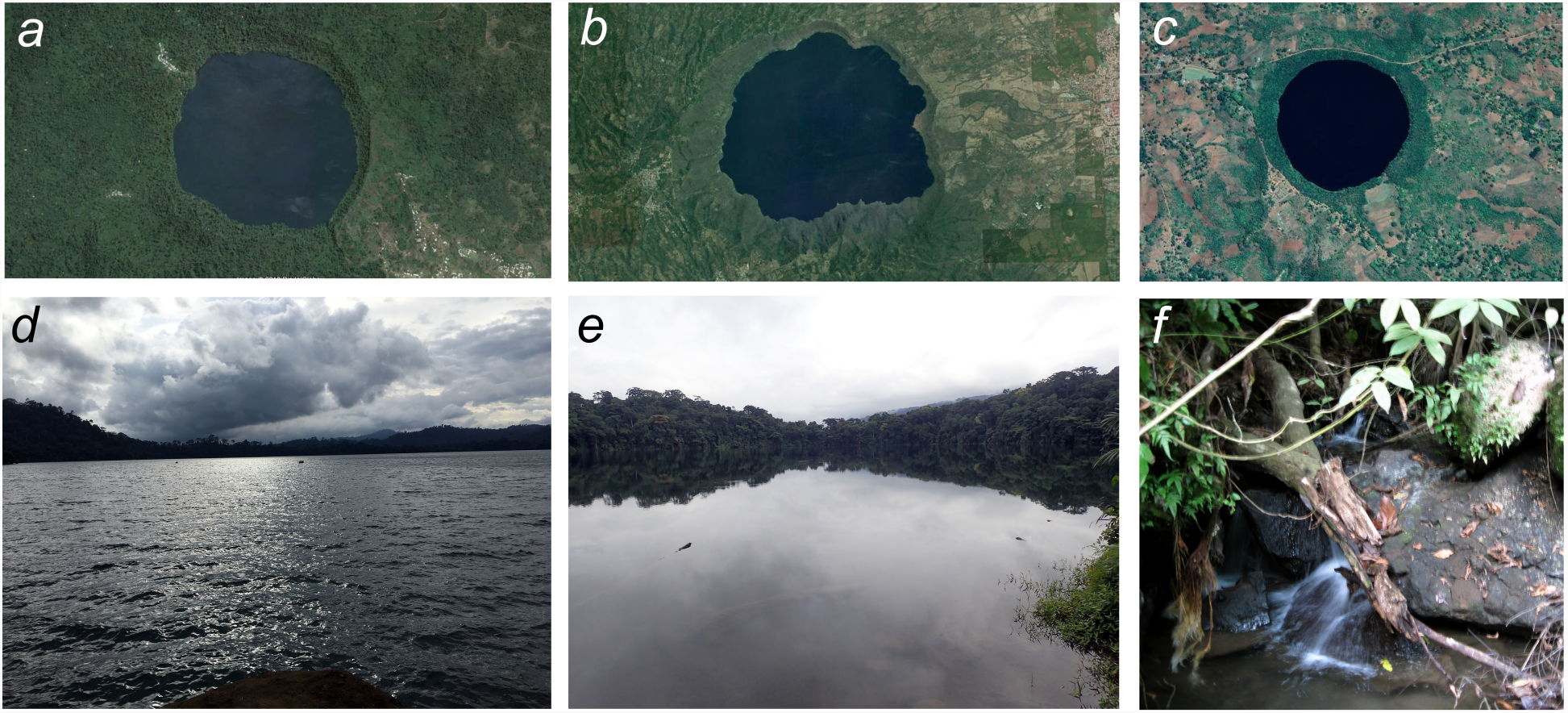
Examples of volcanic crater lakes containing endemic cichlid radiations around the globe: *a,d,f*) Barombi Mbo, Cameroon and its only outlet stream; *b*) Lake Apoyo, Nicaragua, *c*) Lake Massoko, Tanzania, *e*) Lake Bermin, Cameroon. Satellite images (a-c) from Google Earth; (d-f) by CHM.

The evidence for secondary gene flow is remarkably similar across all crater lake cichlid radiations examined with genomic data so far. Admixture proportions with outgroups are frequently detected within the range of 1-4%: 0.6% in Barombi Mbo *Sarotherodon* (percentage of polyphyletic trees in *Saguaro*: [56]), 1.1% in Massoko *Astatotilapia* (Patterson’s *D*: [52]), 4.3% in Apoyo *Amphilophus* (demographic model:[54]), and 4.4% in Ejagham *Coptodon* (1,138 *f*_*D*_ outliers: [98]). No case studies have yet found evidence of substantial divergence in allopatry followed by secondary contact (but see Lake Xiloá *Amphilophus* [54]). Instead, nearly all studies have concluded sympatric divergence with periodic or continuous gene flow, potentially from an initial hybrid swarm population (i.e. introgression from multiple outgroup populations).

Secondary gene flow may have triggered sympatric divergence in a radiation of three *Coptodon* cichlids in Lake Ejagham: demographic analyses of whole genomes suggest that this population did not diversify for 8,000 years despite frequent gene flow until an influx of olfactory receptor alleles 1,000 years ago, coinciding with the first sympatric divergence in the lake [98]. Similarly in Lake Victoria, segregating opsin alleles in riverine cichlid populations were differentially sorted among Lake Victorian cichlids and may have triggered their diversification [32].

Evidence for sympatric divergence in crater lake cichlids without the aid of secondary gene flow remains elusive. Malinsky et al. [52] showed that 1.1% introgression occurred long before the divergence of a shallow/deep-water sister species pair of cichlids in Lake Massoko, Tanzania; however, this initial introgression may have aided later sympatric divergence (which admittedly is very difficult to rule out). Very recent sympatric divergence in some crater lakes or the proliferation of many species from a few colonization events may also suggest that divergence occurred in sympatry without the aid of gene flow [99,100]; however, in the former case it remains unclear if incipient divergence will become stalled as in other sympatric radiations [31]. Very rare secondary gene flow into the Barombi Mbo cichlid radiation (0.6% introgression) without a clear functional role provides weak evidence of sympatric divergence, but more functional characterization and timing of introgression is needed [56]. The recent advent of transgenic reporters, CRISPR-Cas9, and *in situ* hybridization genetic tools within Nicaraguan crater lake cichlids provides much promise for future investigations of the role of introgression in sympatric divergence [81,101].

## *Box 3.* Tools for detecting and timing adaptive introgression

### 1) Detecting and timing introgression

Although there are a variety of tests to detect gene flow on a local scale or within sliding genomic windows, currently three major types of demographic **coalescent** modeling approaches can infer the timing of introgression based on different genomic information: 1) the distribution of allele frequencies from genotype data (site frequency spectrum: e.g; [102,103]), 2) the distribution of haplotype block lengths from phased genomes: e.g; [104,105], and 3) variation in coalescent patterns among gene trees [106].

### 2) Timing of selective sweeps

Recent methods for estimating the age of a selective sweep exploit different aspects about the pattern of variation surrounding the allele on its haplotypic background. These include heuristic approaches that use point estimates of mean haplotype length or the number of derived mutations within a chosen distance of the site [107,108], model-based approaches that use demographic information and summary statistics of allele frequencies and linkage disequilibrium to model a distribution of ages that fit the observed data [109–111], and full sequence approaches that leverage the length of ancestral haplotypes surrounding the beneficial allele and the accumulation of derived mutations [112,113].

### 3) Functional analyses of introgressed variants

Functional annotation of introgressed regions minimally involves searching an annotated reference genome for genes with relevant functions known from model organisms. Intergenic regions can be searched for evidence of strong sequence conservation across taxa [79] or potential regulatory elements (reviewed in [114]). Additionally, genome wide association studies (GWAS) can identify variants in introgressed regions correlated with reproductive isolating barriers. Functional validation of gene and regulatory element variants through genome-editing experiments is also becoming increasingly tractable for non-model organisms (e.g. [81]).

## Glossary Box

### Coalescence

The event of two sampled lineages from different populations merging back in time in a shared ancestral lineage.

### Hybrid swarm

A genetically diverse population with unique allele combinations derived from the hybridization of multiple distinct taxa and subsequent backcrossing with hybrids and crossing between hybrids themselves.

### Introgression

The movement and incorporation of genetic material from one distinct lineage into another upon hybridization between the two and subsequent backcrossing with one of the parent species.

### Linkage disequilibrium

A non-random association of alleles at two or more loci.

### Monophyletic

A group of lineages where the most recent ancestor of the group is not an ancestor of any lineages outside the group.

### Secondary gene flow

Any gene flow event from non-sympatric populations after the initial colonization of the area that sympatric sister species diverged in. Introgression into the diverging sister species following such events potentially brings in variation that has evolved in allopatry that can aid the speciation process.

### Transgressive segregation

The formation of extreme phenotypes in a segregating hybrid population that are outside the range of phenotypes observed in parental species.

## Acknowledgements

We thank A. Foote for helpful comments on an early draft and J. Mallet, D. Matute, A. Meyer, J. McGirr, M. St. John, J. Poelstra, B. Reinhard, and J. Hermisson for valuable discussion of content in this manuscript. This work was supported by the National Science Foundation DEB CAREER Grant #1749764 and by the University of North Carolina to CHM.

## References

1. Mayr, E. (1963) Animal species and evolution, Belknap.

2. Fitzpatrick, B.M. et al. (2009) Pattern, process and geographic modes of speciation. J. Evol. Biol. 22, 2342–2347

3. Fitzpatrick, B.M. et al. (2008) What, if anything, is sympatric speciation? J. Evol. Biol. 21, 1452–9

4. Bolnick, D.I. and Fitzpatrick, B.M. (2007) Sympatric speciation: models and empirical evidence. Annu. Rev. Ecol. Evol. Syst. 38, 459–487

5. Felsenstein, J. (1981) Skepticism towards Santa Rosalia, or why are there so few kinds of animals? Evolution (N. Y). 35, 124–138

6. Dieckmann, U. and Doebeli, M. (1999) On the origin of species by sympatric speciation. Nature 400, 354

7. Gavrilets, S. (2004) Fitness landscape and origin of species, Princeton University Press.

8. Doebeli, M. et al. (2005) What we have also learned: adaptive speciation is theoretically plausible. Evolution (N. Y). 59, 691–699

9. Mallet, J. et al. (2009) Space, sympatry and speciation. J. Evol. Biol. 22, 2332–41

10. Mallet, J. et al. (2009) Space, sympatry and speciation. J. Evol. Biol. 22, 2332–41

11. Foote, A.D. (2018) Sympatric Speciation in the Genomic Era. Trends Ecol. Evol. 33, 85–95

12. Otto, S.P. et al. (2008) Frequency-dependent selection and the evolution of assortative mating. Genetics 179, 2091–2112

13. Coyne, J.A. and Orr, A.H. (1989) Patterns of speciation in Drosophila. Evolution (N. Y). 43, 362–381

14. Polechová, J. et al. (2005) Speciation thrrough competition: a critical review. Evolution (N. Y). 59, 1194–1210

15. Norvaišas, P. and Kisdi, E. (2012) Revisiting Santa Rosalia to Unfold a Degeneracy of Classic Models of Speciation. Am. Nat. 180, 388–393

16. Weissing, F.J. et al. (2011) Adaptive speciation theory: a conceptual review. Behav. Ecol. Sociobiol. 65, 461–480

17. Kopp, M. and Hermisson, J. (2008) Competitive speciation and costs of choosiness. J. Evol. Biol. 21, 1005–1023

18. Nosil, P. and Schluter, D. (2011) The genes underlying the process of speciation. Trends Ecol. Evol. 26, 160–167

19. Ravinet, M. et al. (2017) Interpreting the genomic landscape of speciation: a road map for finding barriers to gene flow. J. Evol. Biol. 30, 1450–1477

20. Matessi, C. et al. (2001) Long-term Buildup of Reproductive Isolation Promoted by Disruptive Selection: How Far Does it Go? Selection 2, 41–64

21. Bolnick, D.I. and Doebeli, M. (2003) Sexual dimorphism: two sides of the same ecological coin. Evolution (N. Y). 57, 2433–2449

22. Bolnick, D.I. (2006) Multi-species outcomes in a common model of sympatric speciation. J. Theor. Biol. 241, 734–744

23. Bürger, R. and Schneider, K.A. (2006) Intraspecific Competitive Divergence and Convergence under Assortative Mating. Am. Nat. 167, 190–205

24. Kirkpatrick, M. and Ravigné, V. (2002) Speciation by Natural and Sexual Selection: Models and Experiments.. Am. Nat. 159, S22–S35

25. Turelli, M. et al. (2001) Theory and speciation. Trends Ecol. Evol. 16, 330–343

26. Garner, A.G. et al. (2018) Genomic signatures of reinforcement. Genes (Basel). 9,

27. van Doorn, G.S. and Weissing, F.J. (2001) Ecological versus Sexual Selection Models of Sympatric Speciation: A Synthesis. Selection 2, 17–40

28. Seehausen, O. (2004) Hybridization and adaptive radiation. Trends Ecol. Evol. 19, 198–207

29. Seehausen, O. (2013) Conditions when hybridization might predispose populations for adaptive radiation. J. Evol. Biol. 26, 279–281

30. Turissini, D.A. and Matute, D.R. (2017) Fine scale mapping of genomic introgressions within the Drosophila yakuba clade. PLOS Genet. 13, e1006971

31. Martin, C.H. (2012) Weak Disruptive Selection and Incomplete Phenotypic Divergence in Two Classic Examples of Sympatric Speciation: Cameroon Crater Lake Cichlids. Am. Nat. 180, E90–E109

32. Meier, J.I. et al. (2017) Ancient hybridization fuels rapid cichlid fish adaptive radiations. Nat. Commun. 8, 14363

33. Wagner, C.E. and McCune, A.R. (2009) Contrasting patterns of spatial genetic structure in sympatric rock-dwelling cichlid fishes. Evolution (N. Y). 63, 1312–1326

34. Servedio, M.R. and Boughman, J.W. (2017) The Role of Sexual Selection in Local Adaptation and Speciation. Annu. Rev. Ecol. Evol. Syst. 48, annurev-ecolsys-110316-022905

35. Kopp, M. et al. (2017) Mechanisms of Assortative Mating in Speciation with Gene Flow: Connecting Theory and Empirical Research. Am. Nat. 191, 1–20

36. Servedio, M.R. et al. (2011) Magic traits in speciation: ‘magic’ but not rare? Trends Ecol. Evol. 26, 389–397

37. Kirkpatrick, M. and Barton, N.H. (2006) Chromosome inversions, local adaptation and speciation. Genetics 173, 419–434

38. Feder, J.L. and Nosil, P. (2009) Chromosomal inversions and species differences: when are genes affecting adaptive divergence and reproductive isolation expected to reside within inversions? Evolution (N. Y). 63, 3061–3075

39. Servedio, M. and Bürger, R. The Effects on Parapatric Divergence of Linkage between Preference and Trait Loci versus Pleiotropy., Genes, 9. (2018)

40. Fuller, Z.L. et al. (2018) Ancestral polymorphisms explain the role of chromosomal inversions in speciation. PLOS Genet. 14, e1007526

41. Gavrilets, S. et al. (2007) Case studies and mathematical models of ecological speciation. 1. Cichlids in a crater lake. Mol. Ecol. 16, 2893–909

42. Gavrilets, S. (2014) Models of speciation: Where are we now? J. Hered. 105, 743–755

43. Coyne, J.A. and Orr, A. (2004) Speciation, Sutherland, MA.

44. Barluenga, M. et al. (2006) Sympatric speciation in Nicaraguan crater lake cichlid fish. Nature 439, 719–723

45. Hahn, M.W. and Nakhleh, L. (2016) Irrational exuberance for resolved species trees. Evolution (N. Y). 70, 7–17

46. Wu, C.-I.I. (2001) The genic view of the process of speciation. J. Evol. Biol. 14, 851–865

47. McGirr, J.A. and Martin, C.H. (2016) Novel candidate genes underlying extreme trophic specialization in Caribbean pupfishes. Mol. Biol. Evol. 34, 873–888

48. Fontaine, M.C. et al. (2015) Extensive introgression in a malaria vector species complex revealed by phylogenomics. Science (80-. ). 347, 1258522–1258522

49. Campbell, C.R. et al. (2018) What is Speciation Genomics? The roles of ecology, gene flow, and genomic architecture in the formation of species. Biol. J. Linn. Soc. at <http://dx.doi.org/10.1093/biolinnean/bly063>

50. Geiger, M.F. et al. (2013) Crater Lake Apoyo revisited: population genetics of an emerging species flock. PLoS One 8, 1–17

51. Igea, J. et al. (2015) A comparative analysis of island floras challenges taxonomy-based biogeographical models of speciation. Evolution (N. Y). 69, 482–491

52. Malinsky, M. et al. (2015) Genomic islands of speciation separate cichlid ecomorphs in an East African crater lake. Science (80-. ). 350, 1493–1498

53. Martin, C.H. et al. (2015) Complex histories of repeated gene flow in Cameroon crater lake cichlids cast doubt on one of the clearest examples of sympatric speciation. Evolution (N. Y). 69, 1406–1422

54. Kautt, A.F. et al. (2016) Multispecies Outcomes of Sympatric Speciation after Admixture with the Source Population in Two Radiations of Nicaraguan Crater Lake Cichlids. PLoS Genet. 12, 1–33

55. Poelstra, J. et al. (2018) Speciation in sympatry with ongoing secondary gene flow and an olfactory trigger in a radiation of Cameroon cichlids. bioRxiv 229864,

56. Richards, E. et al. (2018) Don’t throw out the sympatric speciation with the crater lake water: fine-scale investigation of introgression provides equivocal support for causal role of secondary gene flow in one of the clearest examples of sympatric speciation. bioRxiv at <http://biorxiv.org/content/early/2017/11/17/217984.abstract>

57. Machado-Schiaffino, G. et al. (2015) Parallel evolution in Ugandan crater lakes: Repeated evolution of limnetic body shapes in haplochromine cichlid fish. BMC Evol. Biol. 15,

58. Elmer, K.R. et al. (2014) Parallel evolution of Nicaraguan crater lake cichlid fishes via non-parallel routes. Nat. Commun. 5, 1–8

59. Barluenga, M. and Meyer, A. (2010) Phylogeography, colonization and population history of the Midas cichlid species complex (Amphilophus spp.) in the Nicaraguan crater lakes. BMC Evol. Biol. 10, 326

60. Wu, C.-I. and Ting, C.-T. (2004) Genes and speciation. Nat. Rev. Genet. 5, 114–122

61. Martin, C.H. (2016) The cryptic origins of evolutionary novelty: 1,000-fold-faster trophic diversification rates without increased ecological opportunity or hybrid swarm. Evolution (N. Y). DOI: 10.1111/evo.13046

62. Hedrick, P.W. (2013) Adaptive introgression in animals: Examples and comparison to new mutation and standing variation as sources of adaptive variation. Mol. Ecol. 22, 4606–4618

63. Huerta-Sanchez, E. et al. (2014) Altitude adaptation in Tibetans caused by introgression of Denisovan-like DNA. Nature 512, 194–197

64. Stankowski, S. and Streisfeld, M.A. (2015) Introgressive hybridization facilitates adaptive divergence in a recent radiation of monkeyflowers. Proc. R. Soc. London B 282, 20151666

65. Rieseberg, L.H. et al. (1999) Transgressive segregation, adaptation and speciation. Heredity (Edinb). 83, 363

66. Kagawa, K. and Takimoto, G. (2017) Hybridization can promote adaptive radiation by means of transgressive segregation. Ecol. Lett. 21, 264–274

67. Schumer, M. et al. (2014) How common is homoploid hybrid speciation? Evolution (N. Y). 68, 1553–1560

68. Feder, J.L. et al. (2003) Allopatric genetic origins for sympatric host-plant shifts and race formation in Rhagoletis. Proc. Natl. Acad. Sci. 100, 10314–10319

69. Fishman, L. et al. (2013) Chromosomal rearrangements and the genetics of reproductive barriers in mimulus (monkey flowers). Evolution (N. Y). 67, 2547–2560

70. Guerrero, R.F. and Hahn, M.W. (2017) Speciation as a sieve for ancestral polymorphism. Mol. Ecol. 26, 5362–5368

71. Nelson, T.C. and Cresko, W.A. (2018) Ancient genomic variation underlies repeated ecological adaptation in young stickleback populations. Evol. Lett. 2, 9–21

72. Malinsky, M. et al. (2015) Genomic islands of speciation separate cichlid ecomorphs in an East African crater lake. Science (80-. ). 350, 1493–1498

73. Smith, J.M. and Haigh, J. (1974) The hitch-hiking effect of a favourable gene. Genet. Res. 23, 23–35

74. Nachman, M.W. and Payseur, B.A. (2012) Recombination rate variation and speciation: theoretical predictions and empirical results from rabbits and mice. Philos. Trans. R. Soc. B Biol. Sci. 367, 409–421

75. Cruickshank, T.E. and Hahn, M.W. (2014) Reanalysis suggests that genomic islands of speciation are due to reduced diversity, not reduced gene flow. Mol. Ecol. 23, 3133–3157

76. Stölting, K.N. et al. (2015) Genome-wide patterns of differentiation and spatially varying selection between postglacial recolonization lineages of *Populus alba* (Salicaceae), a widespread forest tree. New Phytol. 207, 723–734

77. Clarkson, C.S. et al. (2014) Adaptive introgression between Anopheles sibling species eliminates a major genomic island but not reproductive isolation. Nat. Commun. 5, 4248

78. Ortíz-Barrientos, D. and Noor, M.A.F. (2005) Evidence for a One-Allele Assortative Mating Locus. Science (80-. ). 310, 1467 LP–1467

79. Siepel, A. et al. (2005) Evolutionarily conserved elements in vertebrate, insect, worm, and yeast genomes. Genome Res. 15, 1034–1050

80. Boyle, E.A. et al. (2017) An Expanded View of Complex Traits: From Polygenic to Omnigenic. Cell 169, 1177–1186

81. Kratochwil, C.F. et al. (2017) Tol2 transposon-mediated transgenesis in the Midas cichlid (Amphilophus citrinellus) — towards understanding gene function and regulatory evolution in an ecological model system for rapid phenotypic diversification. BMC Dev. Biol. 17, 15

82. Cleves, P.A. et al. (2018) An intronic enhancer of Bmp6 underlies evolved tooth gain in sticklebacks. PLOS Genet. 14, e1007449

83. Vijay, N. et al. (2017) Genomewide patterns of variation in genetic diversity are shared among populations, species and higher-order taxa. Mol. Ecol. 26, 4284–4295

84. Van Doren, B.M. et al. (2017) Correlated patterns of genetic diversity and differentiation across an avian family. Mol. Ecol. 26, 3982–3997

85. Ma, T. et al. (2018) Ancient polymorphisms and divergence hitchhiking contribute to genomic islands of divergence within a poplar species complex. Proc. Natl. Acad. Sci. 115, E236 LP–E243

86. Delmore, K.E. et al. (2018) Comparative analysis examining patterns of genomic differentiation across multiple episodes of population divergence in birds. Evol. Lett. 2, 76–87

87. Bush, G.L. (1975) Modes of Animal Speciation. Annu. Rev. Ecol. Syst. 6, 339–364

88. Sorenson, M.D. et al. (2003) Speciation by host switch in brood parasitic indigobirds. Nature 424, 928–931

89. Savolainen, V. et al. (2006) Sympatric speciation in palms on an oceanic island. Nature 441, 210–213

90. Yeaman, S. et al. (2016) The evolution of genomic islands by increased establishment probability of linked alleles. Mol. Ecol. 25, 2542–2558

91. Cotto, O. and Servedio, M.R. (2017) The Roles of Sexual and Viability Selection in the Evolution of Incomplete Reproductive Isolation: From Allopatry to Sympatry. Am. Nat. 190, 680–693

92. Sachdeva, H. and Barton, N.H. (2017) Divergence and evolution of assortative mating in a polygenic trait model of speciation with gene flow. Evolution (N. Y). 71, 1478–1493

93. Seehausen, O. (2006) African cichlid fish: a model system in adaptive radiation research. Proc. R. Soc. Biol. Sci. 273, 1987–98

94. Sparks, J.S. (2004) Molecular phylogeny and biogeography of the Malagasy and South Asian cichlids (Teleostei: Perciformes: Cichlidae). Mol. Phylogenet. Evol. 30, 599–614

95. Schliewen, U.K. et al. (1994) Sympatric speciation suggested by monophyly of crater lake cichlids. Nature 368, 629–631

96. Kautt, A. et al. Multispecies outcomes of sympatric speciation after admixture with the source population in two radiations of Nicaraguan crater lake cichlids. journals.plos.org

97. Wagner, C.E. et al. (2014) Cichlid species-area relationships are shaped by adaptive radiations that scale with area. Ecol. Lett. 17, 583–592

98. Poelstra, J.W. et al. (2018), Speciation in sympatry with ongoing secondary gene flow and a potential olfactory trigger in a radiation of Cameroon cichlids., Molecular Ecology, 27, 1–19

99. Kautt, A.F. et al. (2016) Incipient sympatric speciation in Midas cichlid fish from the youngest and one of the smallest crater lakes in Nicaragua due to differential use of the benthic and limnetic habitats? Ecol. Evol. 6, 5342–5357

100. Elmer, K.R. et al. (2010) Rapid sympatric ecological differentiation of crater lake cichlid fishes within historic times. BMC Biol. 8,

101. Kratochwil, C.F. et al. (2015) Embryonic and larval development in the Midas cichlid fish species flock (Amphilophus spp.): A new evo-devo model for the investigation of adaptive novelties and species differences Evolutionary developmental biology. BMC Dev. Biol. 15, 1–15

102. Excoffier, L. et al. (2013) Robust Demographic Inference from Genomic and SNP Data. PLOS Genet. 9, e1003905

103. Gutenkunst, R.N. et al. (2009) Inferring the Joint Demographic History of Multiple Populations from Multidimensional SNP Frequency Data. PLOS Genet. 5, e1000695

104. Loh, P.-R. et al. (2013) Inferring Admixture Histories of Human Populations Using Linkage Disequilibrium. Genetics 193, 1233 LP–1254

105. Vernot, B. and Akey, J.M. (2014) Resurrecting Surviving Neandertal Lineages from Modern Human Genomes. Science (80-. ). 1245938,

106. Gronau, I. et al. (2011) Bayesian inference of ancient human demography from individual genome sequences. Nat. Genet. 43, 1031

107. Hudson, R.R. (2007) The Variance of Coalescent Time Estimates from DNA Sequences. J. Mol. Evol. 64, 702–705

108. Coop, G. et al. (2008) The Timing of Selection at the Human FOXP2 Gene. Mol. Biol. Evol. 25, 1257–1259

109. Beleza, S. et al. (2013) The Timing of Pigmentation Lightening in Europeans. Mol. Biol. Evol. 30, 24–35

110. Nakagome, S. et al. (2016) Estimating the Ages of Selection Signals from Different Epochs in Human History. Mol. Biol. Evol. 33, 657–669

111. Ormond, L. et al. (2016) Inferring the age of a fixed beneficial allele. Mol. Ecol. 25, 157–169

112. Chen, H. et al. (2015) A hidden Markov model for investigating recent positive selection through haplotype structure. Theor. Popul. Biol. 99, 18–30

113. Smith, J. et al. (2018) Estimating Time to the Common Ancestor for a Beneficial Allele. Mol. Biol. Evol. 35, 1003–1017

114. Chatterjee, S. and Ahituv, N. (2017) Gene Regulatory Elements, Major Drivers of Human Disease. Annu. Rev. Genomics Hum. Genet. 18, 45–63

115. Richards, E.J. and Martin, C.H. (2017) Adaptive introgression from distant Caribbean islands contributed to the diversification of a microendemic adaptive radiation of trophic specialist pupfishes. PLoS Genet. 13, 1–35

